# Prenatal valproic acid-induced autism marmoset model exhibits higher salivary cortisol levels

**DOI:** 10.1101/2022.05.17.492236

**Authors:** Madoka Nakamura, Akiko Nakagami, Keiko Nakagaki, Miyuki Yasue, Nobuyuki Kawai, Noritaka Ichinohe

## Abstract

Individuals with autism spectrum disorder (ASD) are exposed to a variety of stressors owing to their behavioral traits. Cortisol is a hormone typically associated with stress, and its concentration and response to stress are higher in individuals with ASD than in controls. The mechanisms underlying cortisol dysregulation in ASD have been explored in rodents. Although rodent models have successfully replicated the major symptoms of autism (i.e., impaired vocal communication, social interaction deficits, and restricted/repetitive patterns of behavior), evidence suggests that the hypothalamic-pituitary-adrenal (HPA) axis system differs between rodents and primates. We developed an ASD model in the common marmoset (*Callithrix jacchus*), a New World monkey, utilizing prenatal exposure to valproic acid (VPA). In this study, we collected the salivary cortisol levels in VPA-exposed and unexposed marmosets in the morning and afternoon. Our results revealed that both VPA-exposed and unexposed marmosets showed similar diurnal changes in cortisol levels, which were lower in the afternoon than in the morning. However, heightened cortisol levels were observed in VPA-exposed marmosets in the morning, but not in the afternoon. These results are consistent with those of ASD in humans. Our results suggest that VPA-exposed marmosets show similarities not only in their behavioral patterns and brain pathologies, which we have reported previously but also in hormonal regulation, validating the usefulness of VPA-exposed marmosets also as a tool for ASD stress research.

## Introduction

Autism spectrum disorder (ASD) is a neurodevelopmental disorder with a prevalence of 1 in 44 (Maenner et al., 2021). The major symptoms of ASD are impaired vocal communication, deficits in social interaction, and restricted/repetitive behavior patterns (Association and Association, 2013). Individuals with ASD are exposed to a variety of stressors owing to their behavioral traits, and overwhelming stress in ASD can trigger secondary deficits, such as depression and sleep disorders. Cortisol is a hormone typically associated with stress, and its concentration and response to stress are higher in individuals with ASD than in controls. The mechanisms underlying cortisol dysregulation in ASD have been explored in rodent models; however, the hypothalamic-pituitary-adrenal (HPA) axis may differ between rodents and primates (Gibbs, 1986; Broadbear et al., 2004; Goursaud et al., 2006; Bertani et al., 2010). For example, the rodent amygdala is involved in the regulation of basal HPA axis activity, but the primate amygdala is not (Goursaud et al., 2006). Oxytocin, a candidate drug in the treatment of ASD, stimulates adrenocorticotropic hormone release in rats but inhibits its release in primates (Gibbs, 1986). Furthermore, the circadian profiles of the serum hormone levels in rats and primates are different (Bertani et al., 2010). Considering translational research, the primate ASD model appears to be preferable for studying the hormonal stress response in ASD.

We have previously developed an ASD model for a New World monkey, the common marmoset (*Callithrix jacchus*) (Yasue et al., 2015, 2018; Watanabe et al., 2021). The marmosets were prenatally exposed to valproic acid (VPA), which is often used in rodent models of ASD. VPA-exposed marmosets demonstrated all three core symptoms of ASD:1) biased usage of vocal repertoires, 2) weak social attention to unfamiliar conspecifics, and 3) deficits in reversal learning (Watanabe et al., 2021; Nakagami et al., 2022). When inequity between two individuals was introduced (Yasue et al., 2018), VPA-exposed marmosets did not respond negatively even if it was unfavorable to them (i.e., inequity aversion), suggesting a lack of attention toward conspecifics. They also failed to recognize differences between third party reciprocal and non-reciprocal exchanges (6), whereas VPA-unexposed marmosets (UE marmosets) discriminated against these exchanges (Kawai et al., 2014, 2019). Furthermore, transcriptome analyses have revealed that VPA-exposed marmosets replicate a broad range of gene dysregulation in human idiopathic ASD, whereas rodent models generally replicate only a smaller part of the pathology (Watanabe et al., 2021). Thus, VPA-exposed marmosets appear to provide a suitable model for translational research on ASD.

In this study, we collected the salivary cortisol from marmosets in home cage in the morning and afternoon. Salivary collection after adequate training and acclimatization prior to the experiment helped to avoid significant effects on marmoset behavior. We found that VPA-exposed marmosets had heightened basal cortisol levels in the morning, as do humans with ASD. Our results suggest that VPA-exposed marmosets may be useful for translational research on stress pathophysiology in ASD.

## Material and Methods

### Subjects

All experimental animal care procedures were conducted under approved protocols according to the regulations of the National Center of Neurology and Psychiatry (NCNP), Tokyo, Japan. Ten UE marmosets (five males and five females) and nine VPA-exposed marmosets (five males and four females) were included in this study (Table 1). The ages of the experimental animals range from two- to eight-year-old. The mean age was 4.7±1.62 years (UE marmosets:4.5±1.43, VPA-exposed marmosets:4.9±1.79). The subjects were born and raised in a family cage until weaning. During the experimental period, each animal was housed in an individual stainless steel marmoset cage (Natsume Seisakusho Co., Ltd., Tokyo, Japan) at a room temperature of 29±2°C and maintained on a 12h:12h light-dark cycle with ad libitum access to food and water. The lights in the breeding room were turned on at 7:00 a.m. and turned off at 7:00 p.m. daily. Marmosets in the facility were familiar with human contact and approached experimenters to obtain food rewards without hesitation.

**Table 1.** Subject information. 2- to 8-year-old (n=10 males, n=9 females) adult marmosets were used in this study. The mean age of subjects (n=19) in the study was 4.7±1.62 years.

| Animal ID | Age (year) | Sex | Group |
| --- | --- | --- | --- |
| 11111 | 8 | Female | UE |
| 12043 | 7 | Female | VPA |
| 12044 | 7 | Female | VPA |
| 12062 | 7 | Female | VPA |
| 13019 | 6 | Female | UE |
| 14002 | 5 | Female | VPA |
| 14030 | 5 | Male | VPA |
| 14031 | 5 | Male | VPA |
| 14078 | 5 | Male | UE |
| 15033 | 4 | Male | UE |
| 15034 | 4 | Male | UE |
| 15041 | 4 | Female | UE |
| 15049 | 4 | Female | UE |
| 15050 | 4 | Male | UE |
| 16043 | 3 | Male | VPA |
| 16086 | 3 | Female | UE |
| 16089 | 3 | Male | UE |
| 16152 | 3 | Male | VPA |
| 17024 | 2 | Male | VPA |

### VPA Treatment

VPA marmosets were exposed to valproic acid during their fetal stage, whereas UE marmosets were not (Yasue et al., 2015). The dams of VPA-exposed marmosets were housed in their cages. Their blood progesterone levels were monitored periodically to determine the timing of pregnancy, as was done for the UE dams. The VPA group received 200 mg/kg intragastric sodium valproate via an oral catheter daily on days 60 to 66 after conception, for a total of seven treatments. This period was determined based on the administration period (E12 of the rat fetus) used to produce VPA rodent models of ASD. All VPA dams received the medication without vomiting and showed no signs of abnormal pregnancy or delivery. The dams of UE marmosets were administered neither VPA nor a solvent during this period to prevent miscarriage. VPA marmosets displayed no malformations or body weight differences compared with UE marmosets.

### Salivary Collection and Assay

Saliva was sampled to measure cortisol levels twice daily at 7:30 a.m. and at 6:30 p.m., as described in previous research (Kaplan et al., 2012). A thin cotton swab (Matsumotokiyoshi Co., Ltd., Chiba, Japan) was used to collect saliva from the marmosets. Before saliva sampling, the marmosets were fully trained to bite cotton swabs. The cotton swabs were dipped in powdered sugar to ensure that the bites would last sufficiently long (approximately 3-5 min) (Kaplan et al., 2012). A 2.0-mL Coster Spin-X centrifuge tube with a nylon filter (0.22 μm) was filled with the swabs and centrifuged at 10,000 rpm for 5 min to extract the liquid portion of the sample. The collected samples were stored in a −80°C freezer until further use. Saliva samples were collected three times per individual, with no two collections occurring in the same month. Cortisol levels (μg/dl) were measured using an AIA-360 Automated Immunoassay Analyzer with AIA-pack cortisol test cups (Tosoh Corporation, Tokyo, Japan).

## Results

Salivary samples were successfully collected by experimenters while the marmosets were in their home cages. They showed no signs of aggression or aversion during sampling. No significant effect of sex on salivary cortisol levels was observed in both UE- and VPA-exposed groups in the morning and evening. Therefore, we analyzed the data for male and female together.

Figure 1 shows group salivary cortisol levels (mean ± SD) measured in the morning (7:30 a.m.) and afternoon (6:30 p.m.). Both groups exhibited significantly lower cortisol levels in the afternoon than in the morning (p<0.0001 in the UE group, p=0.0001 in the VPA group, Tukey’s HSD test), suggesting that the VPA-exposed marmosets maintained diurnal changes with high cortisol levels in the morning that fell throughout the afternoon. Mean salivary cortisol levels in the VPA-exposed group were significantly higher at 7:30 a.m. than those in the UE group (p=0.0096, Tukey’s HSD test), but there were no significant differences in the afternoon measurements.

**Figure 1.**
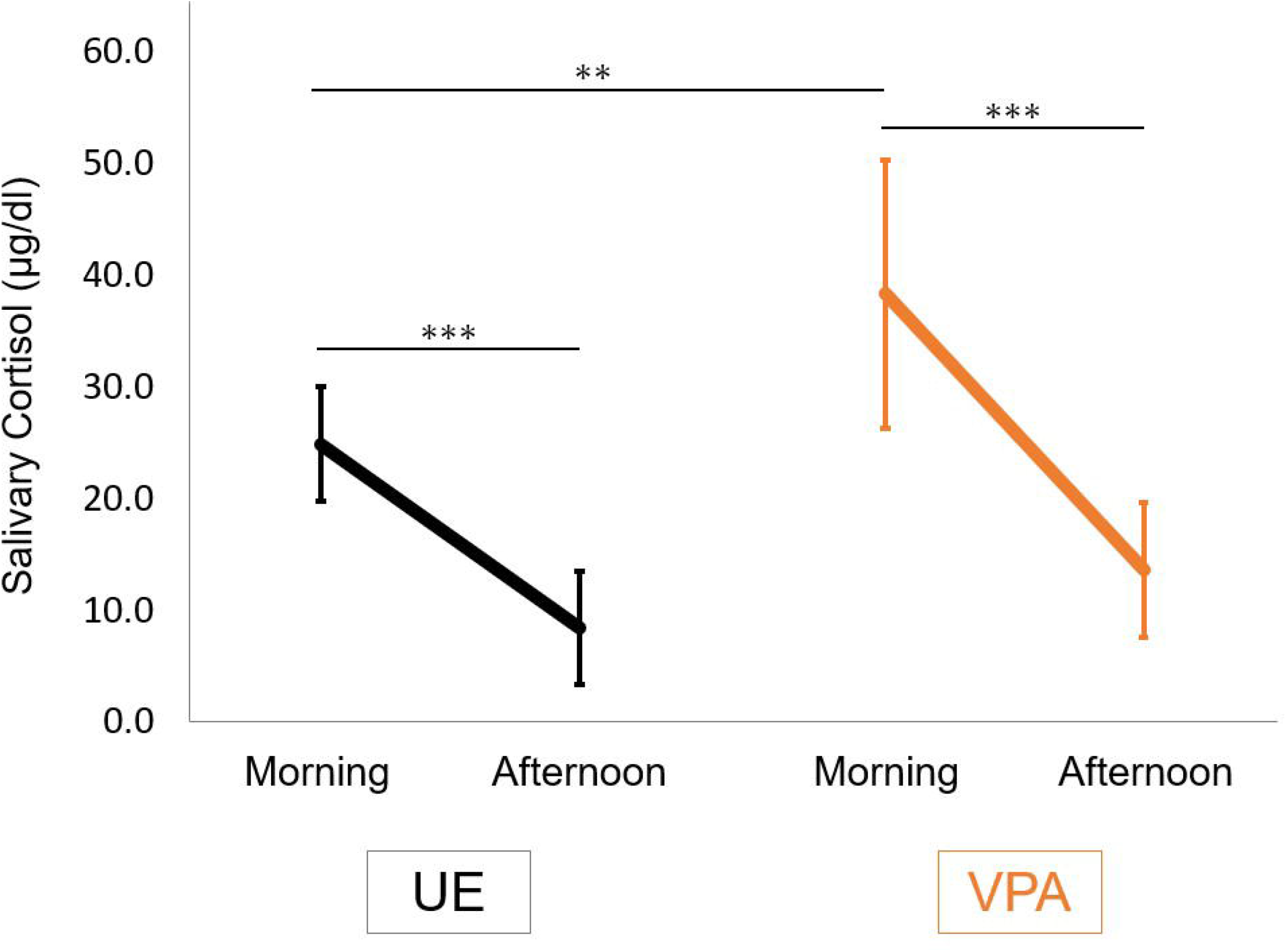
Salivary cortisol levels in the morning (7:30 a.m.) and afternoon (6:30 p.m.) in the valproic acid-exposed (VPA group) and -unexposed (UE group) marmosets. Values are expressed as mean ± standard deviation (n=10 for UE group, n=9 for VPA group, **p<0.01, ***p<0.001).

## Discussion

This study revealed that both UE- and VPA-exposed marmosets showed a similar diurnal change in cortisol levels, which was lower in the afternoon than in the morning. This is consistent with the circadian rhythm of cortisol levels in humans (Smyth et al., 1997). However, heightened cortisol levels in the morning were observed in VPA-exposed marmosets, but not in the afternoon. Previous studies have shown that VPA-exposed marmosets use phee calls more frequently than do UE marmosets (Yamaguchi et al., 2010; Watanabe et al., 2021). This suggests that VPA-exposed marmosets may be in a constant state of stress, such as anxiety. This study is the first case of an ASD model primate showing elevated cortisol.

Studies of cortisol levels in humans with ASD have shown inconsistent results (Tordjman et al., 2014). The discrepancies in their results may be related to study methods (plasma cortisol measures vs. urinary or salivary measures), and sample sizes. We thus compared our results with those of five studies that used a sufficient number of cases with salivary sampling in humans. Four of those reports showed elevated cortisol concentrations (Corbett et al., 2008; Kidd et al., 2012; Spratt et al., 2012; Tordjman et al., 2014). One report commented on the high variability of cortisol concentration in ASD cases (Corbett et al., 2009). In studies examining diurnal variation, two reports observed cortisol increases in the morning and evening (Corbett et al., 2008; Tordjman et al., 2014), while another observed cortisol increases only in the morning (Kidd et al., 2012). Thus, our results are consistent with these reports. VPA-exposed marmosets replicated the abnormal endocrine function observed in people with ASD.

In a rodent model of ASD, BTBR mice and VPA-exposed male rats showed heightened serum cortisol levels (Schneider et al., 2008; Benno et al., 2009; Frye and Llaneza, 2010; Silverman et al., 2010; Ferraro et al., 2021), similar to VPA-exposed marmosets and people with idiopathic ASD. In contrast, transgenic ASD model mice (MeCP2 and FMR1 mutant) showed no elevated levels of cortisol (McGill et al., 2006; Qin and Smith, 2008). It can be difficult, however, to collect specimens for cortisol measurement in rodents without stressing the animals. To avoid this problem, blood is often collected from the heart immediately after euthanasia via acute decapitation. Salivary cortisol collection from rodents also requires restraint and is unsuitable for measuring basal cortisol levels. In this study, we established a method to collect saliva from marmosets after acclimatization by training. The current procedure using marmosets will allow the repeated examination of cortisol levels in ASD models in the same individuals both at basal levels and in the stress response which is also impaired in ASD, and will contribute to a reduction in the number of experimental animals.

This study revealed that VPA-exposed marmosets reproduced the variability of cortisol levels in human ASD. Marmosets are cooperative and highly social primates and are considered a suitable model animal to study stress in social life, and they also show higher social function deficits in cases of ASD (Yasue et al., 2015, 2018; Watanabe et al., 2021; Nakagami et al., 2022). Further examination of cortisol levels in VPA-exposed marmosets would provide a new avenue for studying the biology of stress faced by individuals with ASD and for developing novel therapeutic interventions.

## Acknowledgements

This research was supported by an Intramural Research Grant (grant number 2-7) for Neurological and Psychiatric Disorders from the National Center of Neurology and Psychiatry, by Brain Mapping by Integrated Neurotechnologies for Disease Studies (Brain/MINDS), the Japan Agency for Medical Research and Development (AMED) (22 dm0207066h0004), and by *JPSP* KAKENHI Grant Numbers JP24600020, JP15K01791 to AN, KAKENHI 16H02058, 19K22870, 21H04421, and 21K18552 to NK).

## Author Contributions

N.K., N.I., M.N, and A.N. designed this study. M.N, A.N., and M.Y. performed salivary sampling. M. N. analyzed the data. K.N. managed the production and physical condition of the animals. N. K., N. I. and M. N. wrote the manuscript. All authors have read and approved the final manuscript.

